# Divergent gene expression among phytoplankton taxa in response to upwelling

**DOI:** 10.1101/286138

**Authors:** Robert H. Lampe, Natalie R. Cohen, Kelsey A. Ellis, Kenneth W. Bruland, Maria T. Maldonado, Tawnya D. Peterson, Claire P. Till, Mark A. Brzezinski, Sibel Bargu, Kimberlee Thamatrakoln, Fedor I Kuzminov, Benjamin S. Twining, Adrian Marchetti

## Abstract

Frequent blooms of phytoplankton occur in coastal upwelling zones creating hotspots of biological productivity in the ocean. As cold, nutrient-rich water is brought up to sunlit layers from depth, phytoplankton are also transported upwards to seed surface blooms that are often dominated by diatoms. The physiological response of phytoplankton to this process, commonly referred to as shift-up, is characterized by increases in nitrate assimilation and rapid growth rates. To examine the molecular underpinnings behind this phenomenon, metatranscriptomics was applied to a simulated upwelling experiment using natural phytoplankton communities from the California Upwelling Zone. An increase in diatom growth following five days of incubation was attributed to the genera *Chaetoceros* and *Pseudo-nitzschia*. Here we show that certain bloom-forming diatoms exhibit a distinct transcriptional response that coordinates shift-up where diatoms exhibited the greatest transcriptional change following upwelling; however, comparison of co-expressed genes exposed overrepresentation of distinct sets within each of the dominant phytoplankton groups. The analysis revealed that diatoms frontload genes involved in nitrogen assimilation likely in order to outcompete other groups for available nitrogen during upwelling events. We speculate that the evolutionary success of diatoms may be due, in part, to this proactive response to frequently encountered changes in their environment.

## Introduction

Wind-driven coastal upwelling associated with eastern boundary currents delivers rich supplies of nutrients to illuminated surface waters. This phenomenon provides ideal conditions for blooms of phytoplankton that render coastal upwelling regimes centers of new production even though their relative ocean area is small (Capone and Hutchins, 2013). Typically dominated by large chain-forming diatoms, phytoplankton blooms in upwelling zones rapidly sequester carbon dioxide and are the base of short, efficient food chains that comprise a significant percentage of the global fish catch (Ryther, 1969; Estrada and Blasco, 1985; Lassiter et al., 2006; Lachkar and Gruber, 2013).

The phytoplankton community in upwelling zones is postulated to undergo a ‘conveyer belt cycle’ in which viable cells are upwelled into sunlit waters to seed a surface bloom. The community is then advected away from the upwelled source, and some cells eventually sink out of the photic zone. Surviving cells at depth and positioned in future upwelled waters are able to act as seed stock once winds are favorable for upwelling (Wilkerson and Dugdale, 1987; Wilkerson and Dugdale, 2008). This continuity between a subsurface population and surface bloom during an upwelling event has been observed through a combination of glider and remote sensing techniques (Seegers et al., 2015).

As a result of this vertical transport to a higher light and nutrient-rich environment, phytoplankton exhibit a physiological response, termed shift-up, that includes an acceleration of processes such as nitrate assimilation and growth (MacIsaac et al., 1985; Wilkerson and Dugdale, 1987). An increase in nitrate uptake rates has been repeatedly observed in simulated upwelling mesocosm experiments (Dugdale and Wilkerson, 1989; Fawcett and Ward, 2011) and in a laboratory experiment on the diatom *Skeletonema costatum* (Smith et al., 1992). Shift-up, as expressed through rapid nitrate assimilation, is hypothesized to be linked to the success of diatoms in upwelling regions; it is believed that diatoms respond quickest to available nitrogen once conditions are optimal (Fawcett and Ward, 2011).

Diatoms likely exhibit a unique cellular response that orchestrates this rapid increase in nitrogen assimilation; however, molecular characterization of the shift-up response in general is currently lacking. Only upregulation of the nitrogen assimilation gene, nitrate reductase, has been observed in *Skeletonema costatum* under lab-simulated upwelling conditions, indicating that there is a molecular basis for the shift-up response (Smith et al., 1992). To characterize the phytoplankton community’s response and investigate the molecular basis for shift-up, we applied comparative metatranscriptomics to a simulated upwelling event in a shipboard incubation experiment. Metatranscriptomics is increasingly being utilized with eukaryotic phytoplankton communities to provide a deeper understanding of molecular responses among resident phytoplankton groups (Caron et al., 2017). With the growing availability of reference transcriptomes and genomes of eukaryotic phytoplankton, unprecedented levels and confidence in gene annotation are being obtained from environmental sequences (Keeling et al., 2014; Alexander et al., 2015b). Our results indicate that phytoplankton functional groups exhibit a highly distinct transcriptional response to being upwelled in which diatoms constitutively express genes involved in nitrogen assimilation. This strategy possibly allows diatoms to outcompete other groups for available nitrogen once physical conditions are optimal for growth.

## Results and Discussion

### Experimental overview and physiological observations

Upwelling was simulated by collecting seawater from the 10°C isotherm (96 m) at a site lacking natural upwelling along the California coast (Fig. 1). Satellite-derived sea surface temperature and shipboard wind data indicate that upwelling-favorable conditions were not present for 13 days prior to the incubations, and viable phytoplankton cells were detected at the sampling depth (Supporting Information Fig. S1 and S2). Although multiple factors contribute to the residence time of cells at depth, these data suggest an upper limit of 13 days prior to sampling. Seawater was incubated onboard the ship for up to five days to simulate vertical transport of upwelled waters, although the simulation included a temperature increase that would have likely been more rapid than that of natural upwelling (Supporting Information Fig. S3).

**Fig 1.**
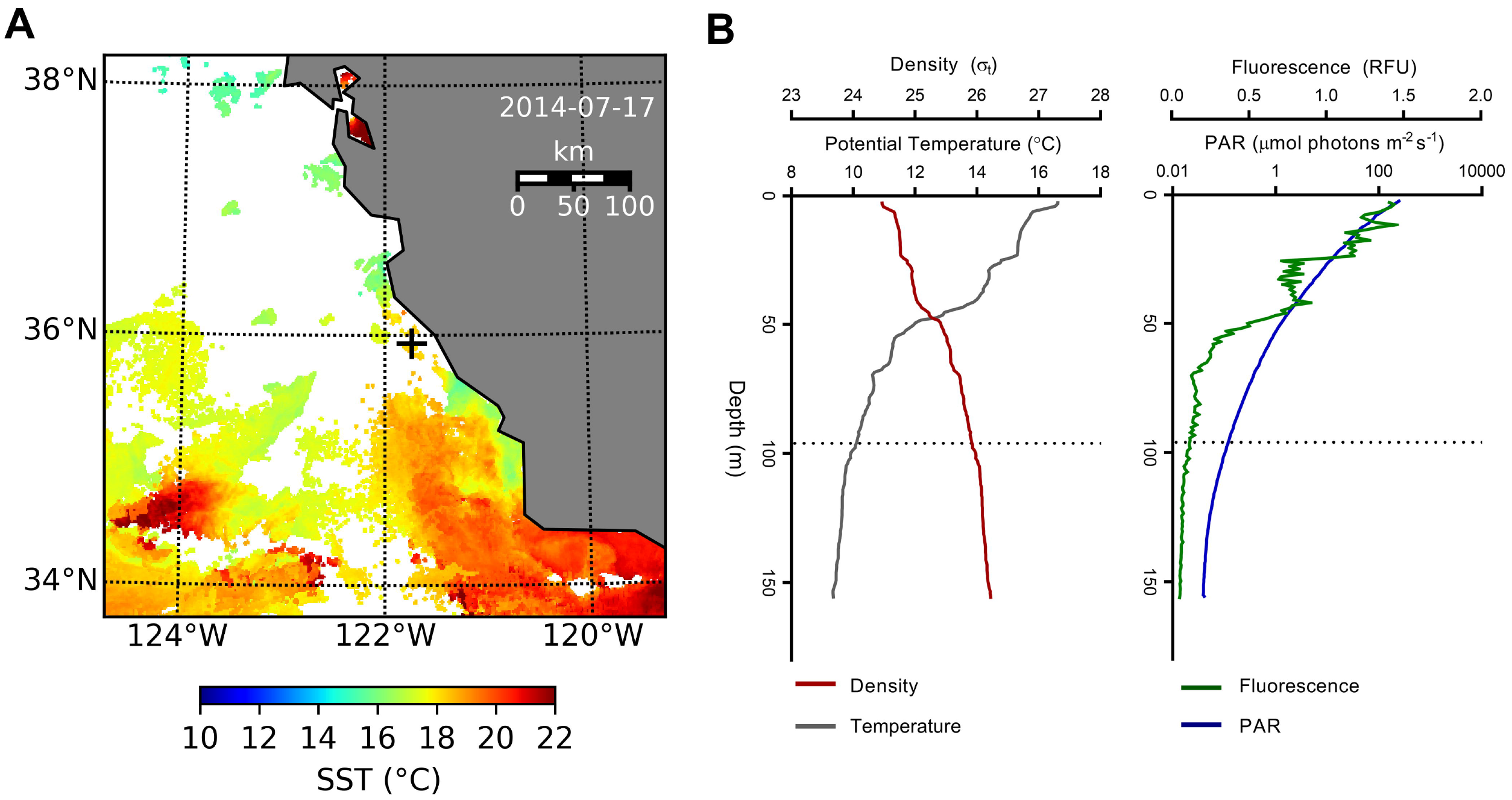
Experimental site (A) Satellite derived sea surface temperature (SST) of the area on the day the incubations began. White areas indicate no data as a result of cloud cover. Water was collected at the point marked with a +. (B) CTD (conductivity-temperature-depth) measurements for potential temperature (°C), density (σ_t_; kg m^−3^), fluorescence (raw fluorescence units), and photosynthetically active radiation (PAR; μmol photons m^−2^ s^−1^) on the afternoon prior to the start of incubations. The dashed line denotes the depth at which water was collected (96 m).

Results from the experiment indicate that a bloom of large phytoplankton (>5 μm) was induced with observations of shift-up, i.e. acceleration, of nitrate uptake and primary production resulting in faster growth rates within these large cells. Macronutrient concentrations in the upwelled waters remained high throughout the incubations; however, significant growth in the large (>5 μm) phytoplankton community was observed (*p* < 0.05; Fig. 2A and Supporting Information Fig. S4A). The initial dissolved iron concentration was approximately 1.28 nmol L^−1^ which is marginally higher than the typical values (<1 nmol L^−1^) observed in the region. For complete drawdown of nitrate, an iron to nitrate ratio of 8 nmol L^−1^:20 μmol L^−1^ is generally required (Bruland et al., 2001). The initial ratio of 1.28 nmol L^−1^:21.86 μmol L^−1^ in the incubations therefore indicates that iron had the potential to be a limiting nutrient which resulted in 15 μmol L^−1^ of unused nitrate after 120 hours.

**Fig 2.**
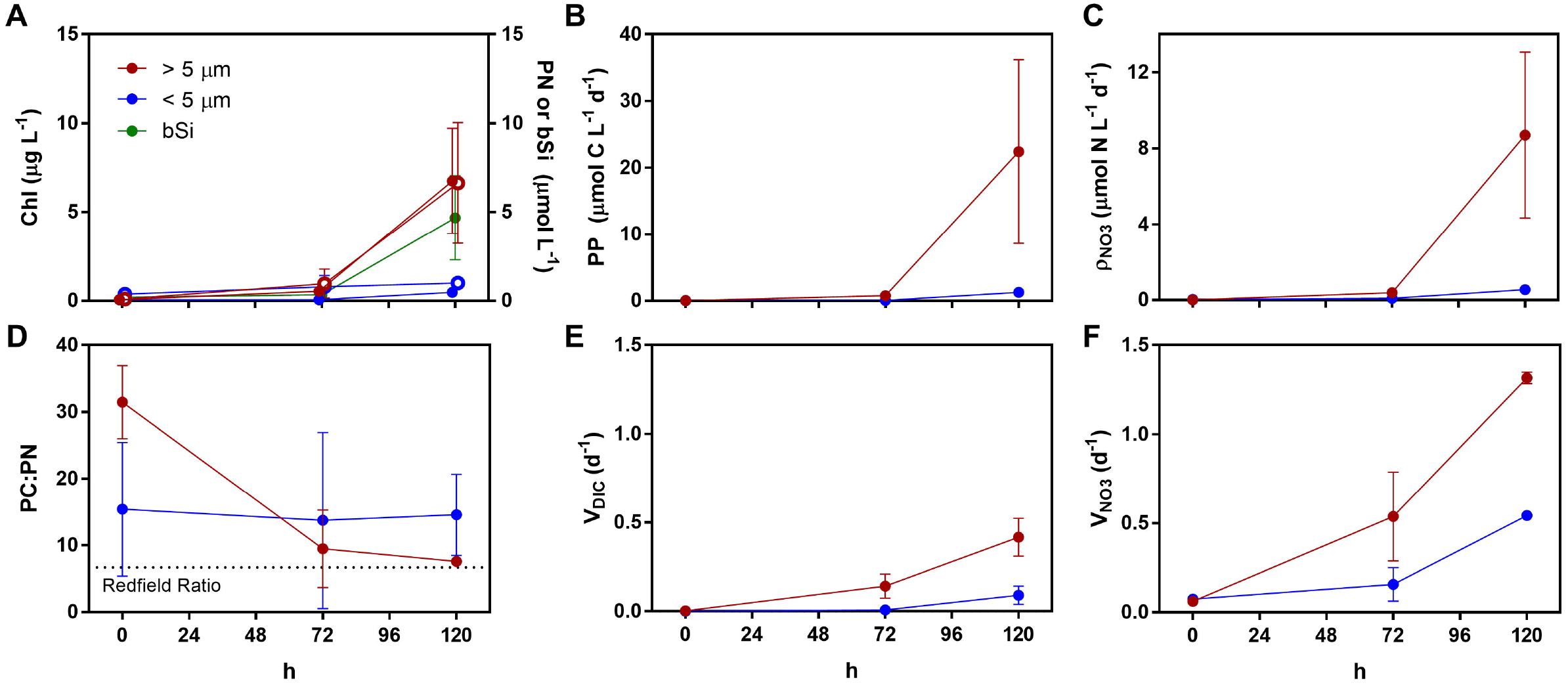
Measurements from the initial upwelled water and incubations at 72 and 120 hours for the community >5 μm (red) and <5 μm (blue). (A) Chlorophyll *a* (closed circles), particulate nitrogen (open circles), and biogenic silica (green). (B) Primary productivity expressed as inorganic carbon uptake (μmol C L^−1^ d^−1^). (C) Absolute nitrate (NO_3_^−^) uptake rates (*ρ*, NO_3_^−^ taken up per unit time). (D) Ratios of particulate carbon to particulate nitrogen. (E) Biomass-specific dissolved inorganic carbon uptake rates (V_DIC_), i.e. inorganic carbon uptake normalized to biomass as particulate carbon (μmol C L^−1^ d^−1^ / μmol C L^−1^ or d^−1^) (F) Biomass-specific nitrate uptake rates (V_NO3_), i.e. nitrate uptake rates normalized to biomass as particulate nitrogen (μmol N L^−1^ d^−1^ / μmol N L^−1^ or d^−1^). Error bars indicate standard deviation of the mean (n = 3).

This success of large phytoplankton is consistent with previous studies showing phytoplankton from large size fractions as the significant contributors to growth and new production during upwelling. Large phytoplankton consistently have greater increases in biomass and appear to outcompete small cells for nutrients during a bloom although reduced grazing pressure may also have an effect (Wilkerson et al., 2000; Fawcett and Ward, 2011). Early on in the incubations, grazers of large phytoplankton may not have been abundant enough to control growth compared to those of small phytoplankton (de Baar et al., 2005).

Nevertheless, the large phytoplankton community also exhibited clear physiological responses to being upwelled. Maximum photochemical yields (F_v_:F_m_) of the whole community increased from 0.25 to 0.51 within the first 72 hours (Supporting Information Fig. S4B). Dissolved inorganic carbon and nitrate uptake in the large cells increased throughout the experiment and was significantly higher than the small cells (Fig. 2B and Fig. 2C). The particulate carbon-to-nitrogen ratio (C:N) was initially 31.5:1 in the >5 μm size fraction but decreased to approach the expected elemental composition of 6.6:1 (Redfield et al., 1963), while C:N values remained fairly constant and above the Redfield ratio for the small size fraction (Fig. 2D). This return to Redfield stoichiometry for the larger phytoplankton cells was coupled with increasing biomass-specific NO_3_^−^ uptake rates (V_NO3_) that were approximately double that of biomass-specific carbon uptake rates (V_DIC_) (Fig. 2E and Fig. 2F).

For the larger phytoplankton, these data indicate a positive response once released from light limitation or a possible resting stage. The initial low F_v_:F_m_ signifies that the community was in a quiescent state but able to return to higher photosynthetic efficiencies within 72 hours. A high initial C:N ratio that approaches the Redfield-predicted value has also been observed in similar mesocosm experiments (Kudela and Dugdale, 2000; Fawcett and Ward, 2011). These studies suggest that the initial high C:N ratio indicates severe N limitation which likely occurred as the phytoplankton cells in aged upwelling water began to sink to depth. Once released from light limitation, the community is able to stabilize with large phytoplankton controlling the total C:N as time progresses. It is also possible that there was C-rich detrital material elevating the initial measurement (Fawcett and Ward, 2011) but acceleration of nitrate uptake, especially in relation to carbon uptake, to drive the phytoplankton community towards balanced growth appears to play a role. The larger cells seem to be able to take advantage of nitrate as conditions become optimal and dominate the community since they uptake nitrate at higher rates than the smaller cells.

### Taxonomic Composition

Large phytoplankton typically dominate blooms during upwelling as observed in this experiment (Estrada and Blasco, 1985; Lassiter et al., 2006). These phytoplankton are typically chain-forming colonial diatoms such as *Chaetoceros* spp. and *Pseudo-nitzschia* spp. In addition to chlorophyll *a*, the upwelling simulation produced significant increases in biogenic silica signifying that the phytoplankton growth can mostly be attributed to diatoms (*p* < 0.05; Fig. 2A). Microscopic cell counts indicated that chlorophytes along with dinoflagellates were most abundant whereas initial cell abundances of diatoms were low. Following incubation, diatoms quickly became the dominant phytoplankton group, particularly those members of the genera *Chaetoceros* and *Pseudo-nitzschia* (Table 1).

**Table 1.** Cell abundances from microscopic counts (10^3^ × cells L^−1^ ± 1 standard deviation of the mean of triplicate samples) of four different phytoplankton groups (diatoms, Bacillariophyceae; dinoflagellates, Dinophyceae; green algae, Chlorophyceae; and haptophytes, Haptophyceae) at the initial timpoint and following 72, and 120 hours of incubation. Diatom abundances are shown for the total diatom assemblage (Total), and for members of the genera *Chaetoceros* and *Pseudo-nitzschia*. bd = below detection limit.

| Class |  | T<sub>0</sub> | T<sub>72</sub> | T<sub>120</sub> |
| --- | --- | --- | --- | --- |
| Bacillariophyceae | Total | $5.8 \pm 2.5$ | $250 \pm 55$ | $2500 \pm 170$ |
| | *Chaetoceros* spp. | $0.44 \pm 0.63$ | $89 \pm 38$ | $980 \pm 760$ |
| | *Pseudo-nitzschia* spp. | $0.5 \pm 0.71$ | $66 \pm 55$ | $1200 \pm 710$ |
| Dinophyceae | | $6.1 \pm 3.8$ | $9.7 \pm 4.6$ | $17 \pm 24$ |
| Chlorophyceae | | $40 \pm 13$ | $250 \pm 45$ | $310 \pm 85$ |
| Haptophyceae | | $0.90 \pm 0.14$ | bd | bd |

Obtaining taxonomically-annotated mRNA read counts also allows for inferences of the relative taxonomic compositions. Metatranscriptome assembly resulted in 3.1 million contigs with levels of annotation similar to previous studies utilizing KEGG and reference transcriptomes from the Marine Microbial Eukaryote Transcriptome Sequencing Project (MMETSP; Supporting Information Table S1)(Keeling et al., 2014; Alexander et al., 2015b; Cohen et al., 2017). These relative abundances of transcripts show that the initial community was relatively diverse, and that dinoflagellates were the dominant phytoplankton group in the pre-upwelled subsurface community (Fig. 3A). Although microscopic cell counts suggest chlorophytes may have been more abundant (Table 1), dinoflagellates may have been exhibiting mixotrophy allowing them to be more transcriptionally active. By 72 and 120 hours following incubation, there was an overwhelming increase in the abundance of mRNA reads attributable to diatoms which is consistent with the bulk measurements, microscope counts, and previous studies (Estrada and Blasco, 1985): diatoms were unequivocally the dominant group within the simulated upwelling event.

**Fig 3.**
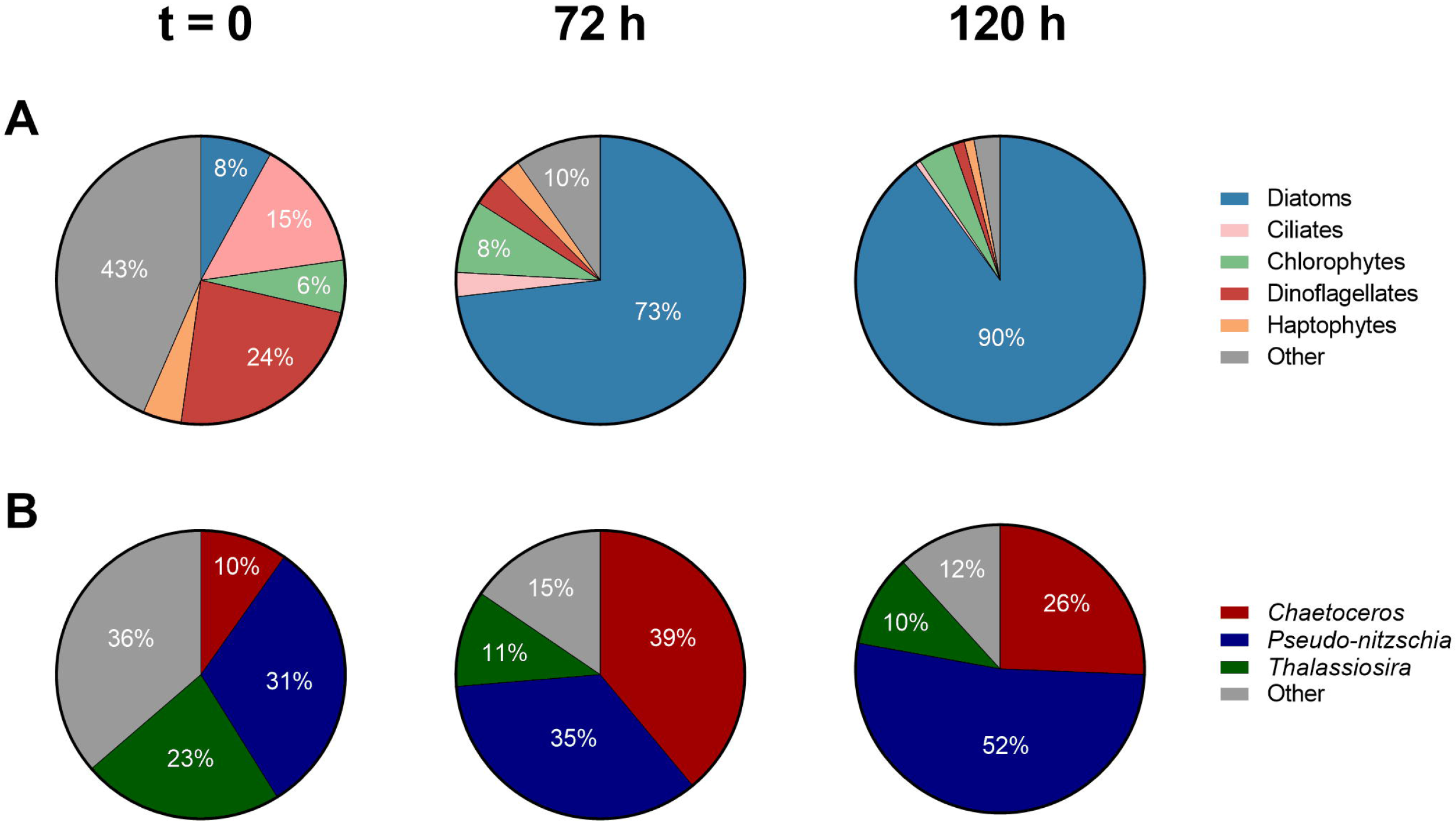
Average taxonomic distribution by mapped reads from each time point. (A) Percentage of reads from the whole community. (B) Percentage of reads for diatom genera within all reads assigned as diatoms.

The taxonomic composition of diatoms followed a similar trend as the whole community with an initially more diverse diatom community that transitioned into one dominated by just two genera: *Chaetoceros* and *Pseudo-nitzschia* (Fig. 3B and Table 1). *Chaetoceros* appeared to make rapid early gains but the community became mostly *Pseudo-nitzschia* by 120 hours. These two genera were also dominant within a previous mesocosm experiment examining shift-up at a nearby coastal California site (Kudela and Dugdale, 2000). *Chaetoceros* spp. were found as resting spores and may quickly germinate following upwelling to make early gains in cell abundance (Pitcher, 1990). Although a resting stage for *Pseudo-nitzschia* spp. is not known (Lelong et al., 2012), they are significant members of the phytoplankton community throughout the upwelling cycle and dominated after 120 hours consistent with a peak in the *Pseudo-nitzschia* produced toxin, domoic acid (Lelong et al., 2012)(Supporting Information Fig. S4C). The average domoic acid concentration was 1.34 μg L^−1^ by 120 hours, and although this concentration is lower than peak concentrations observed during blooms in California coastal waters (Schnetzer et al., 2013), it nevertheless supports that *Pseudo-nitzschia* spp. were abundant within the incubations by 120 hours. This presence of *Pseudo-nitzschia* is unsurprising considering the reports that members of this genus often dominate subsurface chlorophyll maxima (Ryan et al., 2005), thin layers (Rines et al., 2002; McManus et al., 2008), and upwelled communities (Seegers et al., 2015) that often result in harmful algal blooms in this region.

### Comparative Gene Expression of Phytoplankton Groups

Examining shifts in the total transcript pool provides a broad depiction of the responsiveness of different groups. By comparing expression levels at 0 and 72 hours, these shifts reveal the initial whole transcriptome responses to simulated upwelling by the main detected groups of phytoplankton (Fig. 4A). Diatoms had a high proportion of overrepresented genes after upwelling compared to other groups, over 950 (20%) of which were significantly overrepresented (*P*-value < 0.05). Dinoflagellates showed an opposite pattern with gene expression skewed towards overrepresentation in the pre-upwelled condition while haptophytes had an even distribution of overrepresented genes under both conditions. Interestingly, chlorophytes also had a higher number of significantly overrepresented genes post-upwelling, and they were able to maintain their relative proportion of the overall transcript pool unlike the dinoflagellates and haptophytes.

**Fig 4.**
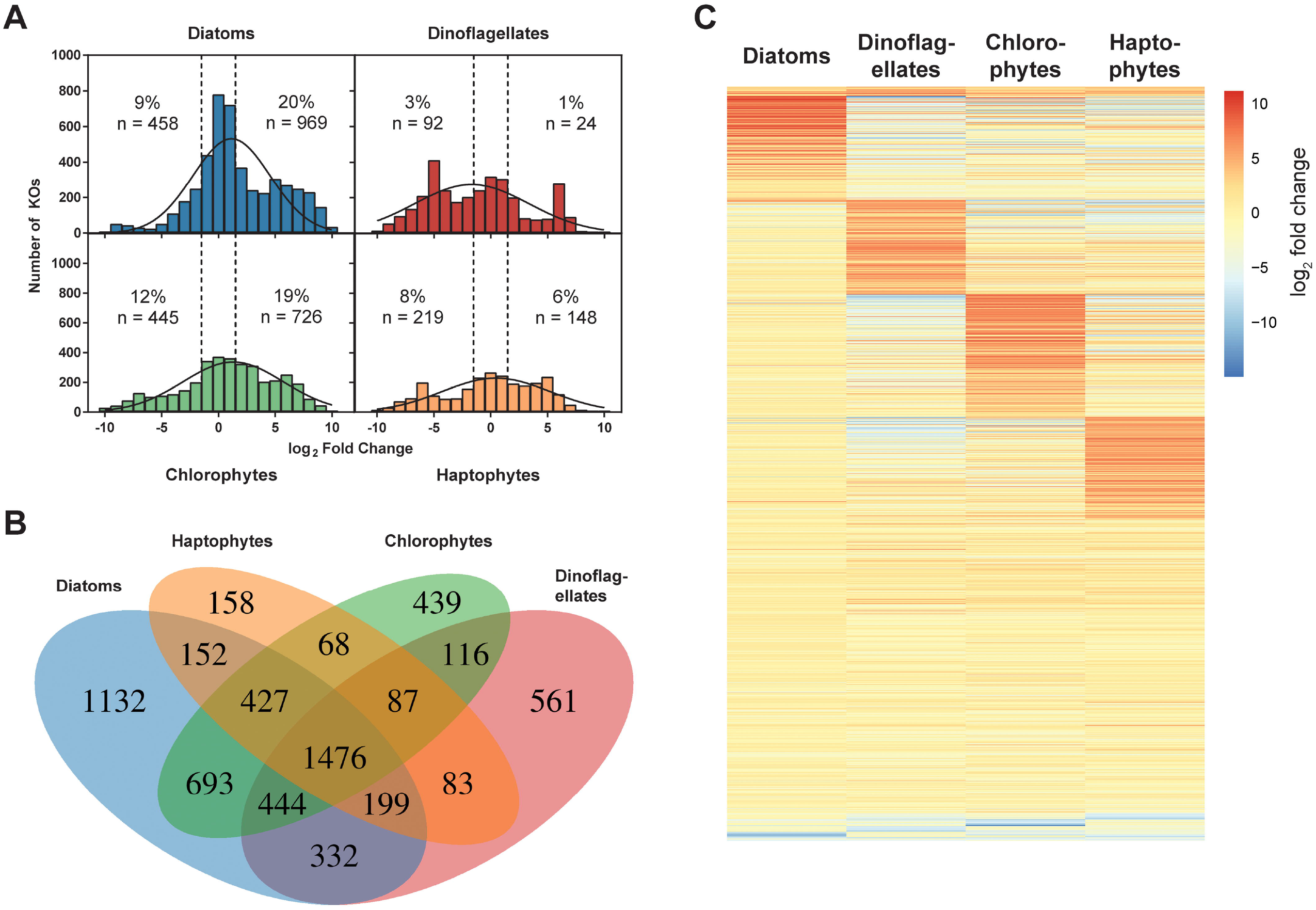
KEGG Ortholog (KO) gene expression comparison among the four main detected phytoplankton groups: diatoms (blue), dinoflagellates (red), chlorophytes (green), and haptophytes (orange). (A) Histograms of KO counts binned by log2 fold change intervals of 1 for 0 and 72 hours and fitted with a Gaussian curve. Dashed vertical lines indicate a log_2_ fold change of −1 or 1. The number and percentage of only the significantly (*P*-value < 0.05) overrepresented genes at 0 hours (pre-upwelling; left) and 72 hours (post-upwelling; right) are annotated on each plot. (B) Venn diagram of expressed KOs at 72 and 0 hours for each group. (C) Heatmap for the 1,476 commonly expressed KOs at 0 and 72 hours. Each row indicates an expressed KO with darker red (positive fold change) indicating overrepresentation at 72 hours and darker blue (negative fold change) indicating overrepresentation at 0 hours.

Smaller changes across all four groups were observed when examining shifts from 72 to 120 hours (Supporting Information Fig. S5). Relatively minor changes in the whole transcript pool and a less pronounced taxonomic shift from 72 to 120 hours indicates that most of the activity in relation to diatom dominance likely occurred in the first 72 hours. This timing and slowing of response also corresponds to field observations that predict a 5-7 day window for cells to achieve balanced growth and transition from shift-up to a low nutrient shift-down (Dugdale and Wilkerson, 1989; Wilkerson et al., 2006). It has been speculated that these shifts, or variable transcript allocation, are a reflection of r- and K-type growth strategies (Alexander et al., 2015b). Our observations appear to follow this paradigm with diatoms exhibiting r-type growth and the highest transcript reallocation in terms of gene count.

Analysis of the expression of genes with shared KEGG Orthology (KO) annotation allows for direct comparisons between taxonomic groups as orthologs normally retain the same function throughout evolutionary history. Similar or different expression of a gene among groups may signify correspondingly similar or different investments in cellular processes at given time points. We detected 1,476 orthologous genes expressed by all four taxonomic groups at 0 or 72 hours (Fig. 4B and Fig. 4C). Only 18 genes were binned as highly overrepresented at 72 hours by all four phytoplankton groups, of which many were related to chlorophyll synthesis. Over 550 genes had low absolute fold change values, many of them positive, across all four groups. These included more photosynthesis-related genes such as photosystem II constituents, photosynthesis electron transport proteins, light-harvesting chlorophyll protein complex proteins, and most of the genes associated with the Calvin cycle. The shared expression of these genes across groups is unsurprising considering the community is transitioning from a deep and dark environment to a sunlit environment, and would benefit from investing in photosynthetic machinery. Other genes that were highly expressed but showed little change in expression across all four groups were associated with other predictable cellular functions such as ribosomal proteins, translation initiation factors, and all constituents of the citric acid cycle.

Of particular interest is the clear overrepresentation at 72 hours of approximately 200 genes per taxonomic group that show little or negative fold change in the other three groups (Fig. 4C and Supporting Information Data Set S1). It is important to note that although differences in shifts in the total transcript pool were observed (Fig, 4A), all groups are still responding and highly increasing their expression of a distinct set of genes compared to the other groups. This pattern continues to hold when examining the genes that were shared between diatoms and just one or two of the other groups (Supporting Information Fig. S6). The genes highly expressed by each group appear to be of diverse function as they do not cluster into certain categories or modules but can be broadly interpreted as investments in different metabolic processes (Supporting Information Fig. S7). These unique responses may reflect fundamental differences in life strategies and ecological traits among functional groups.

To further explain the dominance of diatoms in these systems, expression of diatom annotated genes was investigated. 1,132 KOs were found solely in diatoms, likely due to the abundance of diatoms in our samples resulting in an improved metatranscriptome assembly for that group (Fig. 4B). However, only 173 of these KOs were significantly overrepresented at either 0 or 72 h. It is difficult to determine the importance of the remaining genes as most were expressed in low abundances.

Diatom taxa, however, were not found to respond equivalently to being upwelled; clear differences were noted between *Chaetoceros*, *Pseudo-nitzschia*, and other diatoms (Fig. 5A). Expression of 2,807 orthologs was detected in the genera *Chaetoceros*, *Pseudo-nitzschia*, and all other diatom genera combined mostly consisting of *Thalassiosira*. Similar to what was observed for major taxonomic groups, there was large overrepresentation of distinct sets of genes, particularly in *Chaetoceros* spp., also potentially reflecting transcriptional investments in different processes at different times.

**Fig 5.**
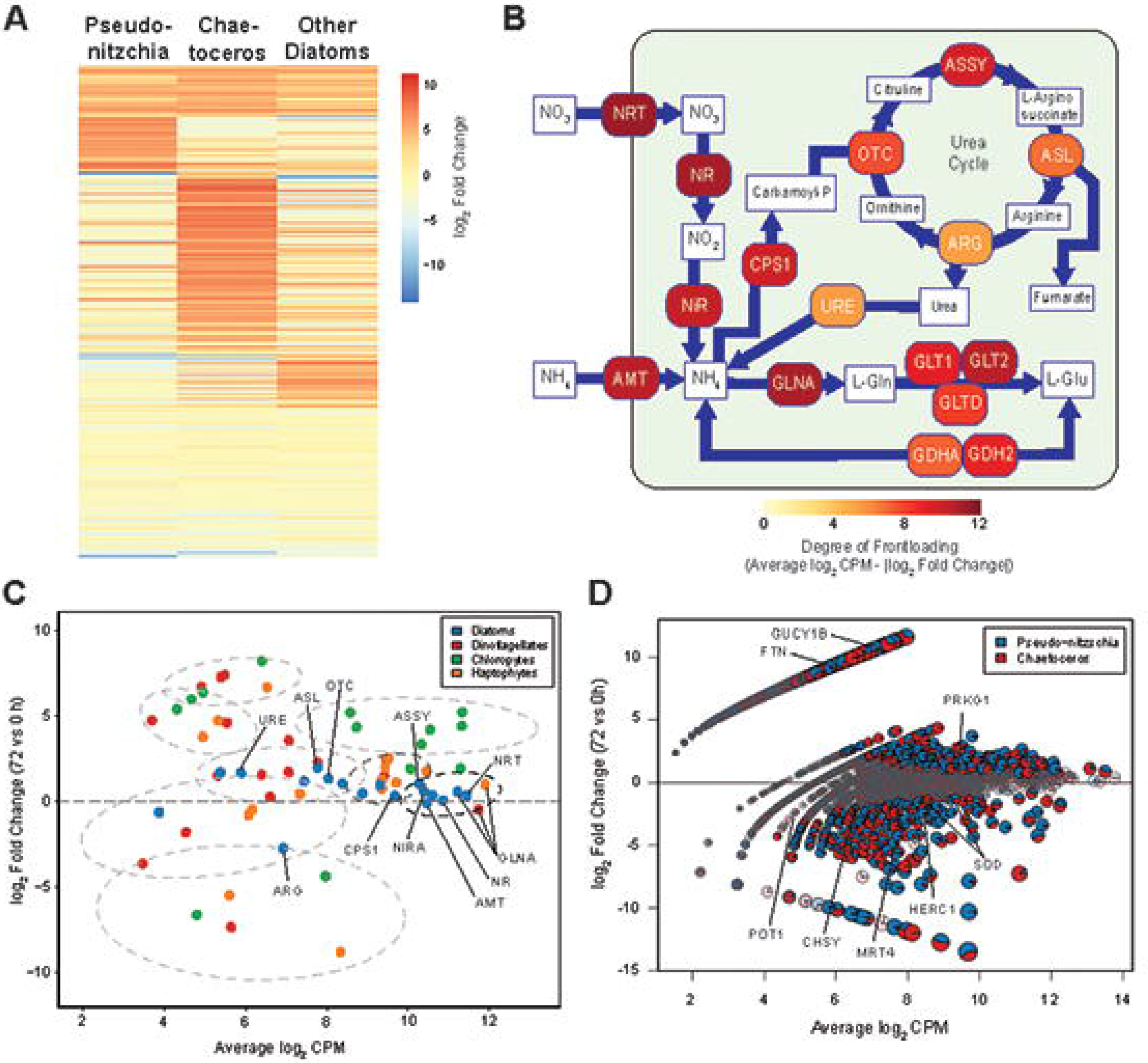
Diatom gene expression. (A) Heatmap for the 2,807 commonly expressed KOs at 0 and 72 hours for *Chaetoceros*, *Pseudo-nitzschia*, and all other diatoms. Each row indicates an expressed KO with darker red (positive fold change) indicating overrepresentation at 72 hours and darker blue (negative fold change) indicating overrepresentation at 0 hours. (B) Cell schematic depicting frontloading of genes associated with nitrogen assimilation and utilization for diatoms. The model is based on Alexander et al. (2015a) and utilizes the same KO numbers. Color indicates the average abundance of the genes (log_2_ CPM) minus the absolute value of the log_2_ fold change to highlight the most abundant, lowest fold change (i.e. frontloaded) genes. Genes are abbreviated as: NRT, nitrate transporter; GLNA, glutamine synthetase; NR, nitrate reductase; AMT, ammonium transporter; NIRA, nitrite reductase (ferredoxin); ASSY, arginosuccinate synthase; CPS, carbamoyl-phosphate synthetase; OTC, ornithine carbamoyltransferase; ASL, argininosuccinate lysase; URE, urease, ARG, arginase; GLT1, glutamate synthase (NADP/NADH); GLT2, glutamate synthase (ferredoxin); GLTD, glutamate synthase; small chain; GDHA/GDH2, glutamate dehydrogenase. (C) MA plot of nitrogen-related genes for the four main detected phytoplankton groups: diatoms (blue), dinoflagellates (red), chlorophytes (green), and haptophytes (orange). Clusters of these N-related genes are shown using k-means clustering (k = 8) with confidence ellipses at the 90% level to aid visualization of frontloading. Select genes are labelled within clusters that represent the most highly frontloaded genes (black ellipse) and additional frontloaded genes (medium gray ellipse). Genes are abbreviated in the same manner as Fig. 4B. (D) Differential transcript abundance between 0 and 72 hours for *Chaetoceros* (red) and *Pseudo-nitzschia* (blue) for expressed KEGG Orthologs (KO). Each pie represents a KO and increases in size with absolute values of its coordinates to optimize visibility. Gene circles that are shaded and have grey borders are not significantly represented in either library (*P*-value ≥ 0.05). Select gene names discussed in the text are labeled as follows: CHSY, chondroitin sulfate synthase; FTN, ferritin; GUCY1B, guanylate cyclase soluble subunit beta; HERC1, ubiquitin-protein ligase; MRT4; mRNA turnover protein, POT1, protection of telomeres protein 1; PRKG1, cGMP-dependent protein kinase 1; SOD, superoxide dismutases.

Ninety-nine significantly differentially expressed genes showed opposing fold-changes in *Chaetoceros* and *Pseudo-nitzschia* when compared to other diatoms (Supporting Information Data Set S2). This highlights that gene expression may not be as accurately assessed by combining genes at high-level taxonomic groupings. The high expression of a gene at one time point or treatment by one group may be cancelled out by another group with opposing expression leading to the incorrect conclusion for the group as a whole. Additionally, one genus could be driving expression of many genes rather than being distributed across the entire group.

### Molecular Characterization of the Nitrogen Assimilation Response

Gene expression was assessed among specific diatom genera and other phytoplankton groups to investigate nitrogen assimilation and utilization, which underlies the shift-up response. Querying nitrogen-related genes for these groups reveals differences in gene expression for the diatoms compared to other phytoplankton (Fig. 5B, Supporting Information Fig. S8). The genes that were most highly expressed both pre- and post-simulated upwelling, referred to here as frontloaded, were almost all from diatoms and related to nitrogen assimilation: nitrate transporter, nitrate reductase, nitrite reductase, and ammonium transporter (Fig. 5C). These frontloaded genes also showed similar expression in diatoms (absolute log_2_ fold change < 1) when comparing the initial and iron-amended treatment following 72 hours, which could be expected as iron was likely not limiting at his time point (Supporting Information Fig. S9).

The only similarly expressed nitrogen-related gene by other groups was glutamine synthetase within dinoflagellates and haptophytes. Within diatoms, the nitrate assimilation genes all had a positive fold change suggesting slightly greater abundance of these genes post-upwelling when compared to pre-upwelling matching our observations of increased nitrate uptake at 72 and 120 hours (Fig. 2C and Fig. 2F). The change in expression in nitrate reductase was very low which contrasts a simulated upwelling experiment with a *Skeletonema* species (Smith et al., 1992). *Skeletonema*, however, was not found to be an abundant genus within this study, and this variation further highlights potential genera-specific differences in response to upwelling.

The urea cycle is believed to facilitate recovery from prolonged nitrogen limitation for diatoms (Allen et al., 2011), but may also be important for the shift-up response. The urea cycle genes carbamoyl-phosphate synthetase and argininosuccinate synthase were also frontloaded by diatoms (Fig. 5B, Fig. 5C, and Supporting Information Fig. S8). Several others were significantly overrepresented post-upwelling including ornithine carbamoyltransferase, arginosuccinate lyase, and urease. The exception in diatoms was arginase, the final enzyme in the urea cycle which was significantly overrepresented pre-upwelling by diatoms not including *Chaetoceros* (Supporting Information Fig. S8). Low expression of arginase post-upwelling is similar to the diatom response to iron enrichment (Marchetti et al., 2012) and may suggest that in both of these scenarios, there are alternative fates for urea cycle intermediates such as nitrogen storage or silica precipitation (Kröger et al., 2001; Llácer et al., 2008).

High relative expression of almost all of these nitrogen-related genes in diatoms compared to most of the other phytoplankton groups at both 0 and 72 hours suggests that investing in nitrogen assimilation and utilization is a priority even when conditions are not optimal for growth. Previous studies indicate that these genes are not constitutively expressed in diatoms, and expression can decline with nitrogen and iron availability (Allen et al., 2008; Allen et al., 2011; Cohen et al., 2017). It is likely that expression of these assimilation genes declines if the community were to face nitrogen limitation at the surface, and at some point at depth, expression returns to high levels.

This transcriptional investment, particularly in the primary genes for nitrate assimilation such as nitrate transporters and nitrate reductase pre-upwelling, may contribute to the rapid response of diatoms as part of their shift-up process. By maintaining elevated pools of nitrogen-related gene transcripts or expressed proteins, upwelled cells are set up to rapidly assimilate available nitrogen whereas other phytoplankton groups only upregulate these genes after being upwelled into the euphotic zone. These results further support the hypothesis that one reason diatoms dominate upwelling regions is because they have the ability to take up and assimilate nitrate more quickly than other phytoplankton groups (Fawcett and Ward, 2011).

Employing this transcriptionally proactive approach to abiotic changes, or ‘frontloading’, has also been characterized with environmental stress response genes in coral and yeast (Berry and Gasch, 2008; Barshis et al., 2013). Additionally, it is similar to what has been observed in diatoms within a previous metatranscriptomic study in relation to iron stress. Iron-enrichment experiments in the northeastern Pacific Ocean demonstrated oceanic diatoms continued expressing genes encoding for iron-free photosynthetic proteins rather than substituting genes encoding for iron-containing functionally equivalent proteins which was different from other phytoplankton groups (Marchetti et al., 2012; Cohen et al., 2017). This strategy is speculated to provide oceanic diatoms with the ability to rapidly acclimate to the inevitable return to iron-limited conditions just as our observations show a strategy that provides certain diatoms with the ability to rapidly take up nitrogen following upwelling. Constitutive frontloading is suggested to provide organisms with resilience to such stressors (Barshis et al., 2013). Along similar lines, bloom-forming diatoms such as *Chaetoceros* and *Pseudo-nitzschia* may have evolved to frontload transcripts of particular genes depending on frequently encountered environmental fluctuations such as conditions associated with the upwelling conveyor belt cycle rather than simply reactively responding to these changes.

### Chaetoceros and Pseudo-nitzschia Expressed Genes

Analyzing genes assigned to two of the most dominant diatom genera, *Chaetoceros* and *Pseudo-nitzschia*, provides further insight into the molecular mechanisms these genera use at depth and as part of their shift-up response. From KOs with module annotations, it is evident that the significantly overrepresented genes at both time points fall into a diverse set of functional categories even at a high level grouping (Supporting Information Fig. S10). To obtain finer resolution, gene expression among all KOs for these genera was examined.

Many overrepresented genes from *Chaetoceros* and *Pseudo-nitzschia* in the pre-upwelled condition indicate a stress response (Fig. 5D). For example, several proteins encoded by highly-expressed genes promote proteasome and ubiquitin activity suggesting that the cells are degrading unneeded or damaged proteins. RNA turnover was also likely increased with high expression of exosome-related transcripts, while expression of protection of telomeres protein promoted stabilization of DNA (Miyoshi et al., 2008). One highly expressed gene was chondroitin sulfate synthase which is potentially related to transparent exopolymer particle (TEP) production (Passow, 2002a; Passow, 2002b). TEP is found to be generated by *Chaetoceros* within the stationary phase which may have contributed to the aggregation and sinking of these cells as well as the high particulate carbon-to-nitrogen ratios observed in the initial community (Fig. 1D).

*Pseudo-nitzchia* in particular expressed a set of distinctive genes as part of its shift-up response. Ferritin was highly expressed post-upwelling, possibly providing a method of storing the essential micronutrient iron (Marchetti et al., 2009). As iron availability in the California upwelling regime can be sporadic and potentially growth limiting, ferritin may provide an advantage to *Pseudo-nitzschia* by concentrating iron for longer-term storage (Bruland et al., 2001) although it may also be used for iron homeostasis (Pfaffen et al., 2015).

At 72 hours, *Pseudo-nitzschia* also uniquely and highly expressed a subunit of soluble guanylate cyclase (sGC or GUCY1B, Fig. 5D). sGC is the only proven receptor of nitric oxide (Denninger and Marletta, 1999) and synthesizes cyclic guanosine monophosphate (cGMP), a second messenger related to many physiological responses (Delledonne, 2005). cGMP activates protein kinase G (PRKG1) which was also significantly expressed (Fig. 5D). Although nitric oxide has been hypothesized to be an infochemical for intercellular signaling and monitoring of stress in diatoms (Vardi, 2008; Amin et al., 2012), *Pseudo-nitzschia* are generally not believed to have a nitric oxide synthase gene as a putative sequence was detected in only one species, *P. multistriata* (Di Dato et al., 2015). *Pseudo-nitzschia* may be utilizing sGC to monitor exhibition of stress from other genera which could allow them to rapidly adapt to changing conditions or respond to sexual cues (Basu et al., 2017). Nitric oxide is also produced by activation of nitrate reductase (Sakihama et al., 2002). As increased nitrate reductase activity occurs as part of the shift-up response, sGC may be used to monitor the continuation of that response and promote certain cellular functions such as gliding of pennates or binary fission (Thompson et al., 2008). Inhibition of sGC prevents the germination of *Leptocyclindrus danicus* resting spores suggesting that it may be involved in transitioning from a resting stage in certain diatoms (Shikata et al., 2011).

Examination of this gene in our reference database reveals that it is highly conserved among *Pseudo-nitzschia* spp. but not ubiquitously present among diatoms (Supporting Information Fig. S11). From this high expression of sGC as part of the upwelling response, evolutionary conservation of this gene, and potential to act as an important signaling device among *Pseudo-nitzschia* spp., we speculate that it may play an important role in shift-up. Additionally, uncharacterized genes that have been previously observed to also increase in expression in comparable microarray studies (Ashworth et al., 2013; Nymark et al., 2013) may elucidate additional mechanisms for the shift-up response in *Chaetoceros* and *Pseudo-nitzschia* (Supporting Information Data Set S3).

### Conclusions

Our simulated upwelling experiment in the California Upwelling Zone is consistent with previous physiological observations of the shift-up response in upwelled phytoplankton: growth of large chain-forming diatoms and increased nitrate assimilation rates. The application of metatranscriptomics to the entire phytoplankton community highlights the divergent transcriptional response of major phytoplankton groups and diatom genera, potentially reflecting variations in their life history strategies. By frontloading nitrogen-related genes, diatoms exhibit the potential for possessing an abundant transcript and/or protein pool allowing them to respond to available nitrate more rapidly than other phytoplankton. This trait is not unlike the response of oceanic diatoms to iron enrichment and may indicate that diatoms have evolved to frontload transcripts in response to frequently encountered changes in their environment. Although the characterization of the shift-up response has largely been focused on nitrogen-related pathways, it is likely that other uncharacterized genes and pathways are also important for diatom success.

## Experimental procedures

### Sample Collection

On 17 July 2014, upwelling conditions were not present at a site within the California Upwelling Zone (35° 56.071’ N, 121° 44.022’ W; Fig. 1A, Supporting Information Fig. S1 and Fig. S2). At 05:00 PDT (12:00 GMT) at the same location, viable phytoplankton cells were detected via imaging flow microscopy (FlowCAM, Fluid Imaging Technologies Inc.) at 96 m although the initial phytoplankton assemblage was small (a total of ~75 cells mL^−1^ greater than 5 μm were found). The sampling depth corresponded to the 10°C isotherm (Fig. 1B).

Seawater from this depth was processed immediately for the initial time point. To simulate upwelling, additional seawater from the same depth was filled into a large acid-rinsed HDPE barrel for homogenization, dispensed to triplicate 10L Cubitainers^®^ (Hedwin Corporation, Newark, DE, USA), and incubated in an on-deck plexiglass incubator with flowthrough seawater at 33% incident irradiance. Seawater collection and incubations followed trace metal clean techniques as they were conducted as part of a larger study to examine diatom responses to iron addition or removal (R. H. Lampe, unpublished). Temperature and on deck irradiance values throughout the incubation are provided in Supporting Information Fig. S3.

Based on macronutrient drawdown, triplicate cubitainers were harvested following 72 hours and 120 hours of incubation. Subsamples from each cubitainer were preserved or measured for chlorophyll *a*, taxonomic composition (by microscopy and FlowCAM), biogenic silica, F_v_:F_m_, domoic acid, nutrients, particulate carbon and nitrogen, carbon and nitrogen uptake, and RNA. Chlorophyll *a*, particulate carbon and nitrogen, and nitrate uptake rates were size fractionated using a series filter cascade while carbon uptake rates were size fractionated using a mesh spacer. Additional methods are described in the Supporting Information.

### CTD, Satellite, and Meteorological Data

Potential temperature, density, photosynthetically active radiation, and fluorescence were obtained from sensors mounted on a 24-bottle rosette onboard the R/V Melville (Seabird 911+ conductivity-temperature depth sensor). Satellite-derived sea surface temperature data on a 0.0125° grid is from the NOAA POES AVHRR satellite courtesy of the NOAA / NESDIS Center for Satellite Applications Research. These data were downloaded from the NOAA CoastWatch Browser and were plotted with matplotlib (Hunter, 2007) for Python v2.7. Wind speed and direction was obtained from the shipboard meteorological system (MetAcq) on the R/V Melville.

### Chlorophyll

Four hundred mL of seawater was gravity-filtered through a 5 μm polycarbonate filter (47 mm) followed by a GF/F filter (25 mm) under gentle vacuum pressure (<100 mm Hg). Filters were rinsed with 0.45 μm filtered seawater and immediately frozen at −80°C until analysis. Chlorophyll *a* extraction was performed using 90% acetone at −20°C for 24 h and measured via *in vitro* fluorometry on a 10-AU fluorometer (Turner Designs, San Jose, CA, USA) using the acidification method (Parsons et al., 1984).

### Biogenic silica

Biogenic silica was determined via filtration of 335 mL onto 1.2 μm polycarbonate filters (45 mm). Concentrations were measured using a NaOH digestion in teflon tubes (Krause et al., 2009) and a colorimetric ammonium molybdate method (Brzezinski and Nelson, 1995).

### Particulate Carbon, Particulate Nitrogen, and Nitrate Uptake

To assess nitrate (NO_3_^−^) uptake rates, 618 mL subsamples were spiked with Na^15^NO_3_ at no more than 10% of ambient nitrate concentration and incubated for eight hours in the flow-through plexiglass incubator. Following incubation, seawater filtration commenced immediately and was performed by gravity through a 5 μm polycarbonate filter (47 mm), and with an in-line vacuum (<100 mm Hg) onto a precombusted (450°C for 5 h) GF/F filter (25 mm). Cells on the 5 μm polycarbonate filter were then rinsed onto an additional precombusted GF/F filter (25 mm) using an artificial saline solution. Filters were then stored at −20°C.

Prior to analysis, filters were dried at 50°C for 24 hours then encapsulated in tin. Particulate nitrogen (PN), particulate carbon (PC), and atom percent of ^15^N were then quantified from this filter using an elemental analyzer paired with an isotope ratio mass spectrometer (EA-IRMS). Samples were not acidified to remove particulate inorganic carbon. Absolute uptake rates (*ρ*, NO_3_ taken up per unit time) were calculated using a constant transport model (Eq. (3) from Dugdale and Wilkerson (1986)). Biomass-specific NO_3_ uptake rates (*V*, NO_3_ taken up per unit PN per unit time) were also calculated according to the constant specific uptake model (Eq. (6) Dugdale and Wilkerson (1986)). ^15^NO_3_ uptake rates were not corrected for possible losses of ^15^N in the form of dissolved organic nitrogen (Bronk et al., 1994); therefore, the reported values are considered conservative estimates or net uptake.

### Dissolved Inorganic Carbon Uptake

Sixty mL samples from each cubitainer were distributed into light and dark bottles cleaned with 1.2 mol L^−1^ HCl. For each bottle, 1.2 μCi of NaH^14^CO_3_ was added and mixed. A 1 mL subsample was taken and added to vials containing 6 mol L^−1^ NaOH to trap and validate the initial inorganic H^14^CO_3_ quantities. The light and dark bottles were incubated on-deck for 6.5-8 h. Samples were filtered onto stacked polycarbonate filters (5 μm and 1 μm) separated with a mesh spacer. Blank control bottles also containing 1.2 μCi of NaH^14^CO_3_ were filtered onto a GF/F filter after 5 min and had counts similar to dark bottles. Filters were vacuumed dried, placed in scintillation vials with 0.5 mL of 6 mol L^−1^ HCl, permitted to degas for 24 h, and counted using a Beckman Coulter LS 6500 scintillation counter. Reported values are light bottles minus dark bottles. Biomass-specific dissolved inorganic carbon (DIC) uptake rates (*V*_*DIC*_) were calculated by normalizing DIC uptake to PC.

### Cell abundances and flow microscopy (FlowCAM)

Phytoplankton cell abundances and species composition were determined by microscopic examination. For microscopic counts, 50 mL samples were preserved in 2% Lugol’s Iodine and settled for >24 hours in Utermöhl chambers (Utermöhl, 1958). Counts were performed at 100x, 200x, and 400x using a Leica DMIL inverted microscope on a minimum of 400 total cells in at least five fields of view.

The viability of cells from the initial seawater and throughout the incubations was monitored through imaging flow microscopy (FlowCAM, Fluid Imaging Technologies Inc., model VS4; prosilica color camera; C71 syringe pump) with Visual Spreadsheets v3.1. The FlowCAM was operated in trigger mode with a 532 nm, 5 mW laser. In this mode, image acquisition by the camera was triggered by chlorophyll or phycoerthrin fluorescence with a minimum threshold of 400 (background noise had a setting of 150-200).

Samples were drawn from the cubitainers into 50 mL Falcon tubes and stored at 4°C in the dark pending processing (typically within 3 hours of collection). At least 5 mL of sample was filtered through a 300 μm nitex mesh screen and passed through the system at a flow rate of 0.2-0.3 mL min^−1^ using a syringe pump equipped with a 5 mL glass syringe. Both a 300 μm flow cell (10x objective) and 100 μm flow cell (20x objective) were used. A digital size filter was applied so that only cells >5 μm were captured in images. The flow cell and tubing were well flushed with Milli-Q water and 70% ethanol between each sample run to avoid cross-contamination.

### RNA Extraction and Sequencing

Seawater was filtered onto 0.8 μm Pall Supor filters (142 mm) using a peristaltic pump, immediately flash frozen in liquid nitrogen, then stored in either liquid nitrogen or at −80°C until extraction. RNA was extracted using the ToTALLY RNA Total RNA Isolation Kit and treated with DNase 1 (Life Technologies, Grand Island, NY, USA). The extraction procedure was followed according to the manufacturer’s instructions with additional first step of glass bead addition to assist with organic matter disruption. RNA quantity and purity was assessed prior to sequencing on an Agilent Bioanalyzer 2100. Total RNA from the triplicate samples for the initial time point (T0) and the first time point (T72) were pooled into one sample due to low RNA yields. Triplicate samples were maintained for the second time point (T120). Library prep was conducted with the Illumina TruSeq Stranded mRNA Library Preparation Kit and HiSeq v4 reagents. Sequencing of barcoded samples was performed on an Illumina HiSeq 2000 (125bp, paired-end).

### Metatranscriptome assembly, annotation, and read quantification

Reads were trimmed for quality and adapter removal using Trimmomatic v0.32 (paired-end mode, adaptive quality trim with 40 bp target length and strictness of 0.6, minimum length of 36 bp)(Bolger et al., 2014). Trimmed paired reads that overlap were merged into single reads with BBMerge v8.0. Merged pairs and non-overlapping paired-end reads from all samples were then used to assemble contigs using ABySS v1.5.2 with varied k-mer sizes (32, 55, 78, and 102)(Birol et al., 2009). Assemblies for each k-mer size were merged using Trans-ABySS v1.5.3 to remove redundant contigs (Robertson et al., 2010), and those shorter than 125 bp were discarded. Read counts were obtained by mapping trimmed reads to contigs with Bowtie2 v2.2.6 (Langmead and Salzberg, 2012) and filtered by mapping quality (MAPQ ≥ 10) with SAMtools v1.2 (Li et al., 2009). Mapping percentages are provided in the Supporting Information Table S3.

Annotation was assigned by best homology (lowest E-value) to protein databases using BLASTX v2.2.31 (E-value ≤ 10^−5^). For taxonomic identification, MarineRefII, a custom reference database was used. MarineRefII contains predicted protein sequences of marine microbial eukaryotes and prokaroytes including all sequenced transcriptomes from the Marine Microbial Eukaryote Transcriptome Sequencing Project (Keeling et al., 2014). MarineRefII was supplemented with transcriptomes of isolated phytoplankton from these incubations adding increased confidence in the taxonomic annotation of some contigs (Supporting Information Table S4). To assign gene function to contigs, the same methodology with the Kyoto Encyclopedia of Genes and Genomes (KEGG; Release 75) was used (Kanehisa et al., 2017). The best hit with a KEGG Ortholog (KO) number from the top 10 hits was chosen. Similarly, analysis of module annotations (MO) was conducted by selecting the top BLASTX hit with a KEGG MO number from the top 10 hits. A summary of annotation results is provided in Supporting Information Table S2.

### Differential expression analysis

Differential expression was assessed by summing read counts of contigs within a taxonomic group (phylum-based or genus for only the diatoms, *Chaetoceros* and *Pseudo-nitzschia*) by KEGG Gene Definition or KEGG Orthology (KO) annotation. edgeR v3.12.0 was used to calculate normalized fold change and counts-per-million (CPM) from pairwise comparisons within each taxonomic group using the exactTest function (Robinson and Smyth, 2008; Robinson et al., 2010). By normalizing within each taxonomic group, shifts in relative abundances are accounted for although reduced sequencing depth for proportionally lower groups may influence fold change estimations for genes detected in one sample but not another. Significance (*P*-value < 0.05) was calculated by using edgeR’s estimate of tagwise dispersions utilizing the available replication within each taxonomic group (Supporting Information Table S2 and Fig. S12)(Chen et al., 2014). ExactTest output in combination with the taxonomic distributions per gene were plotted using a custom plotting function available at https://github.com/marchettilab/mantaPlot.

Shared expression of gene was considered when a gene was detected in at least one of the libraries under comparison for each taxonomic group. For binning of genes displayed in heatmaps (Fig. 4C and Fig. 5A), a positive or negative fold change, variance greater than the number of taxonomic groups, and fold change greater than or less than all other groups were used. Genes with a log_2_ fold change greater than 2 or less than −2 but had a variance less than the number of taxonomic groups were considered similarly overrepresented by all groups. Otherwise, the expression level was considered similar on the basis of fold change. These data were visualized with pheatmap v1.0.8. For KEGG module-based differential expression, quantitative metabolic fingerprinting was used (Alexander et al., 2015a). Briefly, read counts annotated for each KEGG Module category were summed and then normalized by the total number of reads for the time point per functional grouping. These data were also visualized with pheatmap v1.0.8. Gene expression ratios for microarray studies and orthologous genes were obtained from The Diatom Portal (Ashworth et al., 2016).

### Statistical procedures

One-way ANOVAs followed by Dunn’s multiple comparison test were performed on the biological and chemical properties of the seawater (non-gene expression data) in Graphpad PRISM v7.03.

### Data Deposition

The data reported in this paper have been deposited in the National Center for Biotechnology (NCBI) sequence read archive under the accession no. SRP074302 (BioProject accession no. PRJNA320398). Assembled contigs, read counts, and annotations are available on Zenodo (https://doi.org/10.5281/zenodo.1256894). Isolate 18S sequences, transcriptome raw reads, assemblies, and predicted peptide sequences are deposited in Cyverse (http://www.cyverse.org) under the project name unc_phyto_isolates (Supporting Information). Isolate 18S sequences are also deposited in Genbank (accession nos. KX229684-KX229691).

## Acknowledgements

We thank the captain and crew of the R/V Melville and the participants of the IRNBRU cruise: T. Coale (UCSD) performed dissolved nutrient measurements, H. McNair (UCSB) and J. Jones (UCSB) assisted with biogenic silica measurements, M. Maier (OHSU) acquired FlowCAM data, and C. Duckham (UBC) assisted with primary productivity measurements. We also thank W. Burns (UNC), K. Delong (UNC), and C. Payne (UBC) for assistance with sample analyses as well as S. Davies (UNC), W. Gong (UNC), and S. Haines (UNC) for bioinformatic assistance. This work was funded by the National Science Foundation OCE-1334935 (to A.M.), OCE-1334632 (to B.S.T.), OCE-1333929 (to K.T.), OCE-1334387 (to M.A.B.), OCE-1259776 (to K.W.B). Availability of the isolate transcriptomes used in this study was made possible by funding from National Science Foundation OCE-1341479 (to A.M.). R.H.L was partially supported by a fellowship from the UNC Graduate School.

## Supporting Information

**Fig. S1.** Three day average of satellite-derived sea surface temperatures from 02 July 2014 to 4 July 2014. The incubation was started on 16 July 2014 at the site indicated by a +. White areas indicate no data as a result of cloud cover.

**Fig. S2.** Wind speed (m s^−1^) and direction from the R/V Melville for the 48 hours prior to starting the incubations. Spokes display the frequency of winds blowing from particular directions with color-coded bands showing wind speed ranges.

**Fig. S3.** Temperature and on-deck photosynthetically active radiation (PAR) during the incubations. Values for every 15 minutes from a HOBO Data Logger (Onset, Cape Code, MA, USA) are plotted.

**Fig. S4.** Additional measurements from the simulated upwelling incubation experiment conducted in the California Upwelling Zone. (A) Dissolved macronutrient concentrations: nitrate + nitrite (NO_3_^−^ + NO_2_^−^), silicic acid (H_4_SiO_4_), phosphate (PO_4_^3−^), and iron (Fe). All concentrations are in μmol L^−1^ except Fe which is in nmol L^−1^. (B) Particulate domoic acid concentrations. (C) Maximum photochemical yield of photosystem II (F_v_:F_m_). (D) Particulate carbon (PC). PC is the only parameter displayed in this figure with size fractionation: >5 μm (red) and <5 μm (blue). Error bars indicate standard deviation of the mean (n = 3).

**Fig. S5.** Histograms of KEGG Ortholog (KO) gene expression for the four main phytoplankton groups detected in the study. The four main phytoplankton groups are colored as follows: diatoms (blue), dinoflagellates (red), chlorophytes (green), and haptophytes (orange). KO counts are binned by log2 fold change intervals of 1 for 72 and 120 hours and fitted with a Gaussian curve. Dashed vertical lines indicate a log_2_ fold change (120 / 72) of −1 or 1. The number and percentage of significantly overrepresented genes at 120 hours (right) and at 72 hours (left) are annotated on each plot.

**Fig. S6.** Heatmaps for (A) the 1,070 expressed KOs at 72 and 0 h where the gene was not detected in one group other than diatoms. Black bars indicate that the gene was not detected. (B) The 1,170 expressed KOs where the gene was detected in diatoms and only one other group.

**Fig. S7.** Quantitative Metabolic Fingerprint (QMF) depicting relative expression of KEGG modules for each of the four major phytoplankton groups at each time point.

**Fig. S8.** Heatmap of nitrogen related genes for *Chaetoceros*, *Pseudo-nitzschia*, other diatoms, dinoflagellates, chlorophytes, and haptophytes at 0 and 72 hours. The color of each box signifies log_2_ fold change while the numbers in each box denotes the average log_2_ counts-per-million of the gene for that group. Darker red (positive fold change) indicates overrepresentation at 72 hours and darker blue (negative fold change) indicates overrepresentation at 0 hours. Rows are sorted by average abundance among the diatoms.

**Fig. S9.** Fold change of frontloaded nitrogen assimilation genes in diatoms between the iron-amended treatment and control treatment at 72 hours (T72-Fe versus T72-C).

**Fig. S10.** Counts and proportions of significantly overrepresented KEGG orthologs within KEGG module class level two groups at 0 and 72 hours for (A) *Pseudo-nitzschia*, (B) *Chaetoceros*, and (C) other diatoms. None refers to no module annotation. Refer to the Supporting Text for additional discussion.

**Fig. S11.** Phylogenetic tree of guanylate cyclase soluble subunit beta (GUCY1B). The blue branches, k78.1621874 and k78.943480, denote two full length contigs from our metatranscriptome assembly. Bootstrap values ≥ 50 are indicated at the branch points. Refer to the Supporting Text for additional discussion.

**Fig. S12.** Biological coefficient of variation (square root of dispersion) for diatom genes. The biological coefficient of variation is the coefficient of variation with which the true abundance of a gene varies between replicate samples (Chen et al., 2014). The tagwise dispersion based on the trended dispersion was used to assess significance in differential expression.

**Fig. S13.** Phylogenetic tree of nitrate reductase (NR) in diatoms. Contig IDs are colored according to their taxonomic annotation: *Pseudo-nitzschia* (blue) and *Chaetoceros* (red). Branches exclusively ending in *Pseudo-nitzschia* and *Chaetoceros* on the reference tree are also colored correspondingly.

**Table S1**. Summary of functional and taxonomic annotation for the environmental RNA assembly. Functional annotation is from the Kyoto Encyclopedia of Genes and Genomes (KEGG), and taxonomic annotation is from MarineRefII (MRII) supplemented with isolate transcriptomes from the California Upwelling Zone. A definition is assigned when a contig has an acceptable BLASTX hit to the database. Ortholog and module annotation is assigned when a contig has a KEGG ortholog or module annotation from one of the top 10 acceptable BLASTX hits. The top group details contigs with annotations, and the second group details contigs with annotations and reads. Percentages are out of the total number of contigs from assembly: 3,151,426.

**Table S2.** Summary of environmental RNA sequencing with Illumina HiSeq 2000. Samples with (*) denotes samples used in differential expression analysis. All listed samples were used for assembly and estimating dispersions. PE reads and average length in bases were quantified after trimming adapters and for quality from the original 125 base pair reads. Mapped reads are the number and percentage that have a MAPQ score >= 10 from the total number of paired-end (PE) reads

**Table S3.** Summary of transcriptome sequencing, assembly, and gene prediction for isolates from the California Upwelling Zone. Paired-end (PE) reads are the total number of reads after sequencing on an Illumina MiSeq. Average read length given in bases is after trimming the adapters and low quality portions from the original 300 base pair reads. The number of contigs and N50 was generated after assembly with Trinity. Predicted proteins and average protein length in amino acids are derived from GeneMark S-T. 18S sequences, raw sequencing files, assemblies, and predicted peptide sequences are deposited in Cyverse (http://cyverse.org) under the project name unc_phyto_isolates. Genbank accession numbers for 18S sequences are provided in the table.

**Data Set S1.** Uniquely overrepresented genes displayed in Fig. 4C for each taxonomic group.

**Data Set S2.** *Chaetoceros* and *Pseudo-nitzschia* genes that show significantly opposing expression to other diatoms.

**Data Set S3.** Genes of unknown function significantly expressed in our study and with positive fold-changes in similar studies on *T. pseudonana* (tps) and *P. tricornutum* (pti). The study with *T. pseudonana* shows genes with positive fold-changes during exponential growth and light after three days. The dataset with *P. tricornutum* shows genes expressed after 48 of darkness followed by 24 hours of re-exposure to light.

## References

Alexander, H., Jenkins, B.D., Rynearson, T.A., and Dyhrman, S.T. (2015a) Metatranscriptome analyses indicate resource partitioning between diatoms in the field. P Natl Acad Sci USA 112: E2182–E2190.

Alexander, H., Rouco, M., Haley, S.T., Wilson, S.T., Karl, D.M., and Dyhrman, S.T. (2015b) Functional group-specific traits drive phytoplankton dynamics in the oligotrophic ocean. P Natl Acad Sci USA 112: E5972–E5979.

Allen, A.E., Laroche, J., Maheswari, U., Lommer, M., Schauer, N., Lopez, P.J. et al. (2008) Whole-cell response of the pennate diatom Phaeodactylum tricornutum to iron starvation. P Natl Acad Sci USA 105: 10438–10443.

Allen, A.E., Dupont, C.L., Obornik, M., Horak, A., Nunes-Nesi, A., McCrow, J.P. et al. (2011) Evolution and metabolic significance of the urea cycle in photosynthetic diatoms. Nature 473: 203–207.

Amin, S.A., Parker, M.S., and Armbrust, E.V. (2012) Interactions between Diatoms and Bacteria. Microbiol Mol Biol R 76: 667–684.

Ashworth, J., Turkarslan, S., Harris, M., Orellana, M.V., and Baliga, N.S. (2016) Pan-transcriptomic analysis identifies coordinated and orthologous functional modules in the diatoms Thalassiosira pseudonana and Phaeodactylum tricornutum. Mar Genomics 26: 21–28.

Ashworth, J., Coesel, S., Lee, A., Armbrust, E.V., Orellana, M.V., and Baliga, N.S. (2013) Genome-wide diel growth state transitions in the diatom Thalassiosira pseudonana. P Natl Acad Sci USA 110: 7518–7523.

Barshis, D.J., Ladner, J.T., Oliver, T.A., Seneca, F.O., Traylor-Knowles, N., and Palumbi, S.R. (2013) Genomic basis for coral resilience to climate change. P Natl Acad Sci USA 110: 1387–1392.

Basu, S., Patil, S., Mapleson, D., Russo, M.T., Vitale, L., Fevola, C. et al. (2017) Finding a partner in the ocean: molecular and evolutionary bases of the response to sexual cues in a planktonic diatom. New Phytologist 215: 140–156.

Berry, D.B., and Gasch, A.P. (2008) Stress-activated Genomic Expression Changes Serve a Preparative Role for Impending Stress in Yeast. Mol Biol Cell 19: 4580–4587.

Birol, I., Jackman, S.D., Nielsen, C.B., Qian, J.Q., Varhol, R., Stazyk, G. et al. (2009) De novo transcriptome assembly with ABySS. Bioinformatics 25: 2872–2877.

Bolger, A.M., Lohse, M., and Usadel, B. (2014) Trimmomatic: A flexible trimmer for Illumina Sequence Data. Bioinformatics 30: 2114–2120.

Bronk, D.A., Glibert, P.M., and Ward, B.B. (1994) Nitrogen Uptake, Dissolved Organic Nitrogen Release, and New Production. Science 265: 1843–1846.

Bruland, K.W., Rue, E.L., and Smith, G.J. (2001) Iron and macronutrients in California coastal upwelling regimes: Implications for diatom blooms. Limnol Oceanogr 46: 1661–1674.

Brzezinski, M.A., and Nelson, D.M. (1995) The annual silica cycle in the Sargasso Sea near Bermuda. Deep-Sea Res Pt I42: 1215–1237.

Capone, D.G., and Hutchins, D.A. (2013) Microbial biogeochemistry of coastal upwelling regimes in a changing ocean. Nature Geosci 6: 711–717.

Caron, D.A., Alexander, H., Allen, A.E., Archibald, J.M., Armbrust, E.V., Bachy, C. et al. (2017) Probing the evolution, ecology and physiology of marine protists using transcriptomics. Nat Rev Micro 15: 6–20.

Chen, Y., Lun, A., and Smyth, G. (2014) Differential expression analysis of complex RNA-seq experiments using edgeR. In Statistical Analysis of Next Generation Sequence Data. Datta, S., and Nettleton, D.S. (eds). New York: Springer.

Cohen, N.R., Ellis, K.A., Lampe, R.H., McNair, H.M., Twining, B.S., Brzezinski, M.A. et al. (2017) Variations in diatom transcriptional responses to changes in iron availability across ocean provinces. Front Mar Sci 4: 360.

de Baar, H.J.W., Boyd, P.W., Coale, K.H., Landry, M.R., Tsuda, A., Assmy, P. et al. (2005) Synthesis of iron fertilization experiments: From the Iron Age in the Age of Enlightenment. J Geophys Res-Oceans 110: C09S16.

Delledonne, M. (2005) NO news is good news for plants. Curr Opin Plant Biol 8: 390–396.

Denninger, J.W., and Marletta, M.A. (1999) Guanylate cyclase and the ·NO/cGMP signaling pathway. BBA-Bioenergetics 1411: 334–350.

Di Dato, V., Musacchia, F., Petrosino, G., Patil, S., Montresor, M., Sanges, R., and Ferrante, M.I. (2015) Transcriptome sequencing of three Pseudo-nitzschia species reveals comparable gene sets and the presence of Nitric Oxide Synthase genes in diatoms. Sci Rep 5: 12329.

Dugdale, R.C., and Wilkerson, F.P. (1986) The use of 15N to measure nitrogen uptake in eutrophic oceans; experimental considerations. Limnol Oceanogr 31: 673–689.

Dugdale, R.C., and Wilkerson, F.P. (1989) New production in the upwelling center at Point Conception, California: temporal and spatial patterns. Deep-Sea Res 36: 985–1007.

Estrada, M., and Blasco, D. (1985) Phytoplankton assemblages in coastal upwelling areas. In International Symposium on the Most Important Upwelling Areas off Western Africa. Bas, C., Margalef, R., and Rubies, P. (eds). Barcelona, pp. 379–402.

Fawcett, S., and Ward, B. (2011) Phytoplankton succession and nitrogen utilization during the development of an upwelling bloom. Mar Ecol-Prog Ser 428: 13–31.

Hunter, J.D. (2007) Matplotlib: A 2D Graphics Environment. Comput Sci Eng 9: 90–95.

Kanehisa, M., Furumichi, M., Tanabe, M., Sato, Y., and Morishima, K. (2017) KEGG: new perspectives on genomes, pathways, diseases and drugs. Nucleic Acids Res 45: D353–D361.

Keeling, P.J., Burki, F., Wilcox, H.M., Allam, B., Allen, E.E., Amaral-Zettler, L.A. et al. (2014) The Marine Microbial Eukaryote Transcriptome Sequencing Project (MMETSP): Illuminating the Functional Diversity of Eukaryotic Life in the Oceans through Transcriptome Sequencing. PLoS Biol 12: e1001889.

Krause, J.W., Nelson, D.M., and Lomas, M.W. (2009) Biogeochemical responses to late-winter storms in the Sargasso Sea, II: Increased rates of biogenic silica production and export. Deep-Sea Res Pt I 56: 861–874.

Kröger, N., Deutzmann, R., and Sumper, M. (2001) Silica-precipitating Peptides from Diatoms: The Chemical Structure of Silaffin-1A from Cylindrotheca Fusiformis. J Biol Chem 276: 26066–26070.

Kudela, R.M., and Dugdale, R.C. (2000) Nutrient regulation of phytoplankton productivity in Monterey Bay, California. Deep-Sea Res Pt II 47: 1023–1053.

Lachkar, Z., and Gruber, N. (2013) Response of biological production and air-sea CO2 fluxes to upwelling intensification in the California and Canary Current Systems. J Marine Syst 109-110: 149–160.

Langmead, B., and Salzberg, S.L. (2012) Fast gapped-read alignment with Bowtie 2. Nat Meth 9: 357–359.

Lassiter, A.M., Wilkerson, F.P., Dugdale, R.C., and Hogue, V.E. (2006) Phytoplankton assemblages in the CoOP-WEST coastal upwelling area. Deep-Sea Res Pt II 53: 3063–3077.

Lelong, A., Hégaret, H., Soudant, P., and Bates, S.S. (2012) Pseudo-nitzschia (Bacillariophyceae) species, domoic acid and amnesic shellfish poisoning: revisiting previous paradigms. Phycologia 51: 168–216.

Li, H., Handsaker, B., Wysoker, A., Fennell, T., Ruan, J., Homer, N. et al. (2009) The Sequence Alignment/Map format and SAMtools. Bioinformatics 25: 2078–2079.

Llácer, J.L., Fita, I., and Rubio, V. (2008) Arginine and nitrogen storage. Curr Opin Struct Biol 18: 673–681.

MacIsaac, J.J., Dugdale, R.C., Barber, R.T., Blasco, D., and Packard, T.T. (1985) Primary production cycle in an upwelling center. Deep-Sea Res 32: 503–529.

Marchetti, A., Parker, M.S., Moccia, L.P., Lin, E.O., Arrieta, A.L., Ribalet, F. et al. (2009) Ferritin is used for iron storage in bloom-forming marine pennate diatoms. Nature 457: 467–470.

Marchetti, A., Schruth, D.M., Durkin, C.A., Parker, M.S., Kodner, R.B., Berthiaume, C.T. et al. (2012) Comparative metatranscriptomics identifies molecular bases for the physiological responses of phytoplankton to varying iron availability. P Natl Acad Sci USA 109: E317–E325.

McManus, M.A., Kudela, R.M., Silver, M.W., Steward, G.F., Donaghay, P.L., and Sullivan, J.M. (2008) Cryptic Blooms: Are Thin Layers the Missing Connection? Estuar Coast 31: 396–401.

Miyoshi, T., Kanoh, J., Saito, M., and Ishikawa, F. (2008) Fission Yeast Pot1-Tpp1 Protects Telomeres and Regulates Telomere Length. Science 320: 1341–1344.

Nymark, M., Valle, K.C., Hancke, K., Winge, P., Andresen, K., Johnsen, G. et al. (2013) Molecular and Photosynthetic Responses to Prolonged Darkness and Subsequent Acclimation to Re-Illumination in the Diatom Phaeodactylum tricornutum. PLoS ONE 8: e58722.

Parsons, T.R., Maita, Y., and Lalli, C.M. (1984) In A Manual of Chemical & Biological Methods for Seawater Analysis. Amsterdam: Pergamon.

Passow, U. (2002a) Transparent exopolymer particles (TEP) in aquatic environments. Prog Oceanogr 55: 287–333.

Passow, U. (2002b) Production of transparent exopolymer particles (TEP) by phyto-and bacterioplankton. Mar Ecol-Prog Ser 236: 1–12.

Pfaffen, S., Bradley, J.M., Abdulqadir, R., Firme, M.R., Moore, G.R., Le Brun, N.E., and Murphy, M.E.P. (2015) A Diatom Ferritin Optimized for Iron Oxidation but not Iron Storage. J Biol Chem.

Pitcher, G.C. (1990) Phytoplankton seed populations of the Cape Peninsula upwelling plume, with particular reference to resting spores of Chaetoceros (bacillariophyceae) and their role in seeding upwelling waters. Estuar Coast Shelf S31: 283–301.

Redfield, A.C., Ketchum, B.H., and Richards, F.A. (1963) The influence of organisms on the composition of sea water. In The Sea. Hill, M. (ed). New York: Interscience, pp. 26–77.

Rines, J.E.B., Donaghay, P.L., Dekshenieks, M.M., Sullivan, J.M., and Twardowski, M.S. (2002) Thin layers and camouflage: hidden Pseudo-nitzschia spp. (Bacillariophyceae) populations in a fjord in the San Juan Islands, Washington, USA. Mar Ecol-Prog Ser 225: 123–137.

Robertson, G., Schein, J., Chiu, R., Corbett, R., Field, M., Jackman, S.D. et al. (2010) De novo assembly and analysis of RNA-seq data. Nat Meth 7: 909–912.

Robinson, M.D., and Smyth, G.K. (2008) Small-sample estimation of negative binomial dispersion, with applications to SAGE data. Biostatistics 9: 321–332.

Robinson, M.D., McCarthy, D.J., and Smyth, G.K. (2010) edgeR: a Bioconductor package for differential expression analysis of digital gene expression data. Bioinformatics 26: 139–140.

Ryan, J.P., Chavez, F.P., and Bellingham, J.G. (2005) Physical-biological coupling in Monterey Bay, California: topographic influences on phytoplankton ecology. Mar Ecol-Prog Ser 287: 23–32.

Ryther, J.H. (1969) Photosynthesis and Fish Production in the Sea. Science 166: 72–76.

Sakihama, Y., Nakamura, S., and Yamasaki, H. (2002) Nitric Oxide Production Mediated by Nitrate Reductase in the Green Alga Chlamydomonas reinhardtii: an Alternative NO Production Pathway in Photosynthetic Organisms. Plant Cell Physiol 43: 290–297.

Schnetzer, A., Jones, B.H., Schaffner, R.A., Cetinic, I., Fitzpatrick, E., Miller, P.E. et al. (2013) Coastal upwelling linked to toxic Pseudo-nitzschia australis blooms in Los Angeles coastal waters, 2005-2007. J Plankton Res 35: 1080–1092.

Seegers, B.N., Birch, J.M., Marin, R., Scholin, C.A., Caron, D.A., Seubert, E.L. et al. (2015) Subsurface seeding of surface harmful algal blooms observed through the integration of autonomous gliders, moored environmental sample processors, and satellite remote sensing in southern California. Limnol Oceanogr 60: 754–764.

Shikata, T., Iseki, M., Matsunaga, S., Higashi, S.-i., Kamei, Y., and Watanabe, M. (2011) Blue and Red Light-Induced Germination of Resting Spores in the Red-Tide Diatom Leptocylindrus danicus†. Photochem Photobiol 87: 590–597.

Smith, G.J., Zimmerman, R.C., and Alberte, R.S. (1992) Molecular and physiological responses of diatoms to variable levels of irradiance and nitrogen availability: growth of Skeletonema costatum in simulated upwelling conditions. Limnol Oceanogr 37: 989–1007.

Thompson, S.E.M., Taylor, A.R., Brownlee, C., Callow, M.E., and Callow, J.A. (2008) The Role of Nitric Oxdie in Diatom Adhesion in Relation to Substratum Properties. J Phycol 44: 967–976.

Utermöhl, H. (1958) Zur Vervollkommnung der quantitativen Phytoplankton-Methodik. Mitt int Ver theor angew Limnol 9: 1–38.

Vardi, A. (2008) Cell signaling in marine diatoms. Commun Integr Biol 1: 134–136.

Wilkerson, F.P., and Dugdale, R.C. (1987) The Use of Large Shipboard Barrels and Drifters to Study the Effects of Coastal Upwelling on Phytoplankton Dynamics. Limnol Oceanogr 32: 368–382.

Wilkerson, F.P., and Dugdale, R.C. (2008) Nitrogen in the Marine Environment (2nd Edition). Burlington, MA, USA: Academic Press.

Wilkerson, F.P., Dugdale, R.C., Kudela, R.M., and Chavez, F.P. (2000) Biomass and productivity in Monterey Bay, California: contribution of the large phytoplankton. Deep-Sea Res Pt II 47: 1003–1022.

Wilkerson, F.P., Lassiter, A.M., Dugdale, R.C., Marchi, A., and Hogue, V.E. (2006) The phytoplankton bloom response to wind events and upwelled nutrients during the CoOP WEST study. Deep-Sea Research Part Ii-Topical Studies in Oceanography 53: 3023–3048.

